# Computational study of *Piper betle* L. phytocompounds by *insilico and ADMET* analysis for prediction of potential xanthine oxidase inhibitory activity

**DOI:** 10.1101/2023.01.13.523909

**Authors:** P. Vikrama Chakravarthi, M. Karthikeyan, L. Lakshmanan, S. Murugesan, A. Arivuchelvan, K. Sukumar, A. Arulmozhi, A. Jagadeeswaran

**Affiliations:** Department of Veterinary Pharmacology and Toxicology, Veterinary College and Research Institute, Tamil Nadu Veterinary and Animal Sciences University, Namakkal, India

**Keywords:** *P.betle*, GC-MS, *Insilico* study & ADMET analysis

## Abstract

**Background:** Xanthine oxidase (XO) enzyme is directly associated with pathogenesis of gout and indirectly with cancer, diabetes and metabolic syndromes. Allopurinol is a xanthine oxidase inhibitor which is useful in the treatment of gout but it causes side effects in humans and birds. Therefore, this study sought to identify the alternative compound from the natural resources with fewer side effects than the conventional ones.

**Objectives:** The present study was designed to findout the xanthine oxidase inhibitory activity of *Piper betle* phytocompounds in comparison with Allopurinol.

**Methods:** The detection of phytocompounds present in the *P. betle* L. extract was identified through Gas Chromatography Mass Spectrometry (GC-MS) analysis. The *insilico* analysis and ADMET properties were evaluated for the GC-MS derived *P.betle* phytocompounds. Among the 32 phyto compounds of *P.betle*, 18 were showed favourable affinity with xanthine oxidase and their different conformational structures were docked in schrodinger module with xanthine oxidase enzyme structure. The interpretation of results was carried out through GLIDE XP Scoring, GLIDE Energy, MM-GBSA energy and hydrogen bond and Pi-Pi interactions. Further the absorption, distribution, metabolism, excretion and toxicity (ADMET) analysis were also carriedout using QIKPRO programme.

**Results:** The results revealed that, the twelve phytocompounds of *P.betle* showed GLIDE XP docking score (−6.881 to - 4.766 Kcal/mol) higher than allopurinol (−4.535 Kcal/mol) and 11 phytocompounds showed higher GLIDE energy (−42.822 Kcal/mol to - 36.706 Kcal/mol) than allopurinol (−32.676 Kcal/mol). Chromonol and eugenol were the most potential compounds of *P.betle* which showed both hydrogen bond and Pi-Pi interactions with the target xanthine oxidase as that of standard drug allopurinol. The selected phytocompounds satisfied the ADME descriptors and have no violation of Lipinski’s rule of five.

**Conclusions:** The *insilico* and ADMET profile study on the phytocompounds of *P.betle* predicted their promising potential xanthine oxidase inhibition activity and they could be developed as alternate molecules against synthetic agents by *invivo* experiments for the treatment of xanthine oxidase associated diseases like hyperuricimia and cardiovascular disorders.

## Introduction

Xanthine oxidase (xanthine oxidoreductase - XOR) is an evolutionarily conserved housekeeping enzyme with a principal role in purine catabolism (Hille and Nishino, 1995). It catalyzes the hydroxylation of hypoxanthine (HX) to xanthine and then to uric acid with simultaneous production of reactive oxygen species (ROS). In most living species, such as primates, this metabolic pathway has been highly conserved during the evolutionary process, while in birds and Dalmatian dogs have experienced an increase in circulating levels of uric acid after losing the functionality of the final step in the degradation of uric acid (Oda et al., 2002). Humans with mutations in XOR suffer from xanthinuria, an autosomal recessive disorder resulting in kidney stone formation and urinary tract disorders (Ichida *et al*. 1997).

A number of studies have revealed that uric acid is an independent risk factor for cardiometabolic diseases as shown in figure 1, suggesting that uric acid may be a potential therapeutic target for the treatment of hyperuricemia associated diseases (Lee *et al*., 2020). In addition, xanthine oxidase enzyme is also involved in the production of hydrogen peroxides and superoxide anions as byproducts during the formation of uric acid which subsequently increases the cellular oxidative stress (Nile and Khobragade, 2011). The oxidative stress is related to endothelial dysfunction and ischemia-reperfusion injury, and may be implicated in the pathogenesis of heart failure, hypertension, and ischemic heart disease. Since, the hyperuricimia is not only a risk factor for gout, it also recognized to increase the chance of developing cardiovascular disease, type 2 diabetes mellitus, and Chronic kidney disease in humans (Galassi and Borghi, 2015). The experimental evidence suggests that hyperuricaemia may also play a causal role in the etiopathogenesis of insulin resistance (a known cardiovascular risk factor) through the induction of systemic endothelial dysfunction and pro-inflammatory and oxidative alterations, especially at the level of adipocytes (Sautin, 2007).

**Figure 1.**
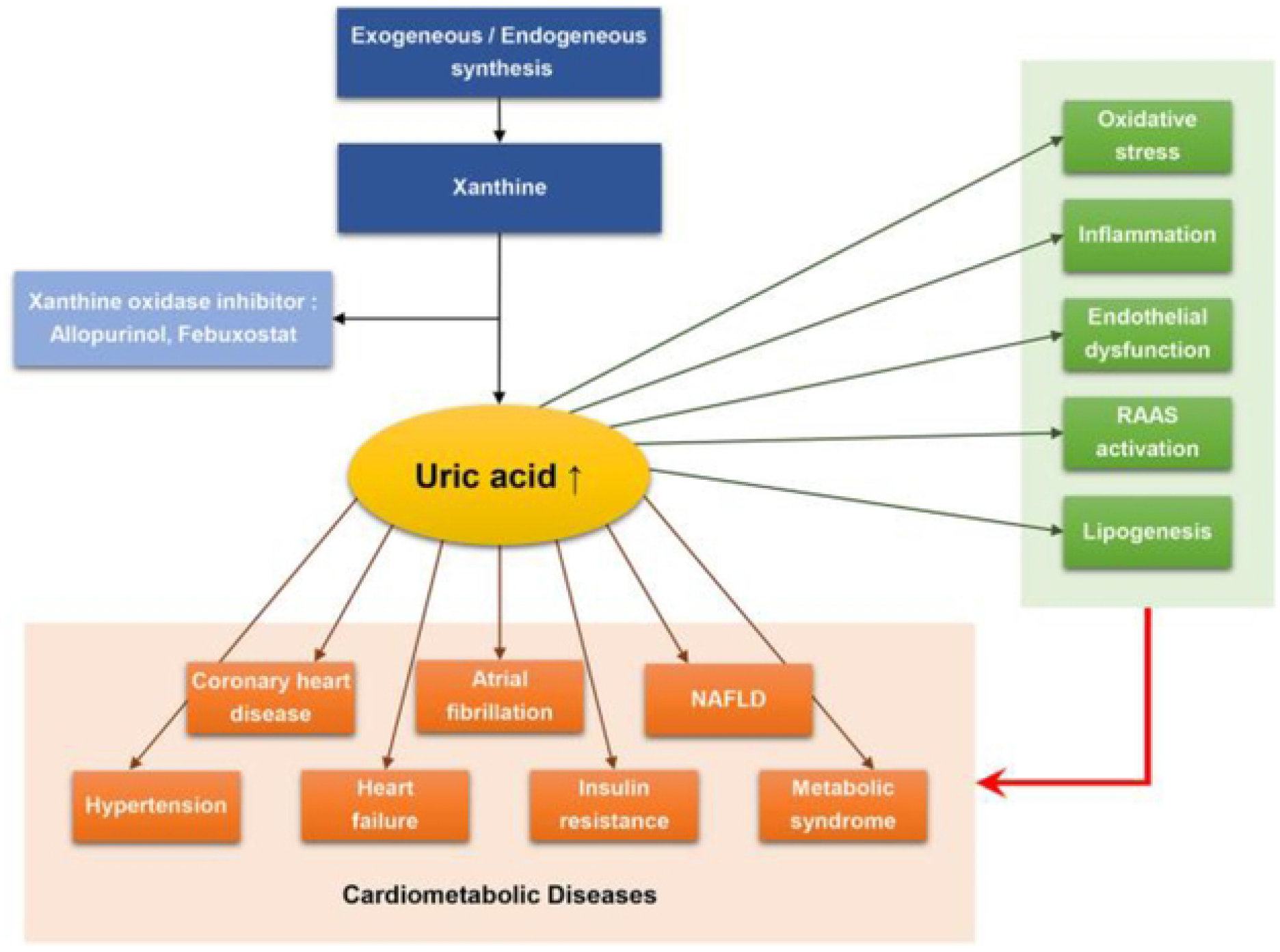
Schematic diagram showing interplay of uric acid, metabolic syndrome and cardiovascular disease RAS - renin-angiotensin aldosterone system; NAFLD - Non-alcoholic fatty liver disease (Lee et al., 2020)

Xanthine oxidase inhibitors (XOIs) are perhaps showed the most positive effects on cardiovascular outcomes beyond their treatment effect on gout (Mercuro *et al*. 2004). Allopurinol is commonly used for the treatment of gout to suppress uric acid generation (Borges *et al*., 2002, Chen *et al*., 2016). However, the drug use is limited by the occurrence of side effects like fatal liver necrosis (Pereira *et al*., 1998), renal toxicity (Horiuchi *et al*., 2000) and hypersensitivity problems (Hammer *et al*., 2001) in humans. Also, the residual toxicity by the use of allopurinol in the liver tissue was reported in broiler chicken (Settle *et al*., 2012).

Hence to circumvent such problems efforts have been made to identify compounds with xanthine oxidase inhibitory activities from natural sources (Kong *et al*., 2000). New herbal inhibitors of xanthine oxidase discovered by using molecular modeling techniques such as molecular docking and molecular dynamics simulations. The combinatorial chemistry and high-throughput screening have increased the possibility of finding new lead compounds at much shorter time periods than conventional medicinal chemistry. However, too much promising drug candidates often fail because of unsatisfactory absorption, distribution, metabolism, excretion and toxicity (ADMET) properties. *In silico* ADMET studies are expected to reduce the risk of late-stage attrition of drug development and to optimize screening and testing by looking at only the promising compounds. Several methods of integrating ADME properties to predict pharmacokinetics at the organ or body level have been studied (Yamashita and Hashita, 2004). QikProp is one of the well-known absorption, distribution, metabolism, excretion and toxicity (ADMET) prediction program (Jorgensen and Duffy, 2002) which could be used to get reliable results.

Therefore, this study has been designed to identify a potential xanthine oxidase enzyme inhibitor from natural products with fewer side effects than those of conventional drugs for treating gout using computational approaches. The plant derived molecule based *in silico* virtual screening is an efficient method currently used for drug development research. Using these methods, it is possible to find new lead molecules quickly at a moderate cost (Kikiowo et al., 2020). Among the available herbs, *Piper betle* L. was an important species of the Piperaceae family with glossy heart-shaped leaves which acts as magnificent reservoirs of phenolic compounds. Herbs containing phytocompounds, such as polyphenols and flavonoids, have been used as xanthine oxidase inhibitors (Costantino et al. 1996). Phytochemical investigations on *P.betle* leaves revealed the presence of alkaloids, flavonoids, phenolic compounds and terpenoids (Vikrama Chakravarthi, *et al*. 2022). Experimentally *P.betle* leaves have been found to reveal diverse pharmacological actions, such as antimicrobial (Shitut et al., 1999), hepatoprotective (Saravanan et al., 2002) antioxidant and antiinflammatory (Pin et al., 2010) effects. With those backgrounds, the present study was conducted to identify the xanthine oxidase inhibitory activity of *P.betle* phytocompounds.

## Materials and Methods

The local variety (Karpoori) of *P. betle* L. herb was collected from different regions of Namakkal District of South India, Tamil Nadu and authenticated by the Botanical Survey of India (No.BSI/SRI/5/23/2017/Tech/1921), Southern Regional Centre, Government of India, Coimbatore, Tamil Nadu. The freshly collected leaves were shade dried to prepare the herbal extract. The dried leaves were finely powdered using a mechanical mixer and collected in clean polythene bags. Alcoholic extract was prepared using 100 grams of *P. betle* L. powder in 400 ml of ethanol. The mixture was agitated in an orbital shaker at 200 RPM for 72 h and evaporated at 35°C under reduced pressure using a rotary evaporator and preserved for phytochemical analysis. Gas Chromatography-Mass Spectrometry (GC-MS) analysis was performed to identify the bioactive components present in the extract. The analysis was performed in GC-MS 5975 C Agilent System and Turbo Mass software was adopted to handle mass spectra and chromatograms (Adams, 2007).

### *In silico Docking* study

*In silico* docking studies was performed with Schrodinger Masetro software using Glide module for enzyme ligand docking (Friesner *et al*., 2004). The study was conducted in the Computer Aided Drug Design (CADD) laboratory of department of bioinformatics, Alagappa university, Karaikudi, Tamil Nadu,India.

### Modelling platform

The entire computational analysis was carried out using the Schrodinger-Maestro software packages (10.2 version tool) including ligand preparation, GLIDE (Grid-based Ligand Docking with Energetics) docking, grid generation and free energy calculations.

### Target protein preparation

Xanthine oxidase model from bovine milk source was downloaded from the Research Collaboratory for Structural Bioinformatics (RCSB) protein data bank. Prior to docking of ligands (phytochemicals/drug) into the protein’s (xanthine oxidase) active site (allopurinol binding site), the protein was prepared using protein preparation wizard of Schrodinger’s molecular docking software.

### Ligand preparation

A total of 32 phytocompounds from *P. betle* derived through GC-MS analysis were used for molecular docking studies. Allopurinol structure was retrieved from drug bank. All the ligand molecules were prepared using LigPrep 2.4 and the selected ligands were sketched in Marvin Sketch (Freeware) and saved as SDF format for docking with target protein.

### Molecular docking

All the ligand molecules were docked into the binding site of xanthine oxidase using the GLIDE based ligand docking program of Schrodinger software. Searches were made for favorable docking interactions between one or more ligand molecules (Phytochemicals / drug) with target protein (xanthine oxidase). The docking calculations were generated few poses for each ligand with target protein. The selection of the best pose was done based on the interaction between the ligand and the target protein.

### GLIDE Scoring, GLIDE Energy and MM-GBSA Energy

GLIDE score is an empirical scoring function designed to maximize separation of compounds with strong binding affinity from those with little to no binding ability. GLIDE XP (Extra precision) score was based on ligand-protein docked complex contacts, hydrogen bonds and Pi-Pi (benzene ring) interactions and varderwalls forces. The scoring of compounds was used to rank the ligand-protein docked complex. The GLIDE energy was derived based on the electrostatic interaction of docked complex within atom level. Also, in the present study the Molecular Mechanics Generalized Born Surface Area (MM-GBSA) module was used to estimate the free energy of binding between ligand and target protein (Subramaniyan *et al*., 2018). The total free energy binding was estimated as follows using the software:

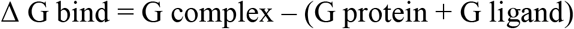

Where each energy term is a combination of G = Molecular mechanics energies (MME) + GSGB (SGB solvation model for polar solvation) + GNP (non-polar solvation) (Kakarala and Jamil, 2015) coulomb energy, covalent binding energy, Vander Waals energy, lipophilic energy, GB electrostatic solvation energy, prime energy, hydrogen-bonding energy, hydrophobic contact, and self-contact correction (Lee and Olson, 2013).

### Absorption, distribution, metabolism, excretion and toxicity (ADMET) prediction

Absorption, distribution, metabolism, excretion and toxicity (ADMET) analysis of molecules was performed for the selected six phytocompounds of *P.betle* (4-chromanal, Allyl diacetoxy benzene, Allyl 6 methoxyphenol, Methyl octadecadienol, Eugenol and Benzoic acid 3,3 dimethyl) from the results of induced fit docking study using QikProp program of Schrödinger suite (Jorgensen *et al*., 2002). QikProp program (QikProp, version 3.5, Schrödinger, LLC, New York, NY, 2012) generates set of physicochemically significant descriptors which further evaluates ADMET properties. The descriptors include molecular weight, H bond acceptor, H bond donor, QPlogPo/w, QPlogS, QPPCaco, QPlogBB and Rule of Five.

The drug likeness of the six *P.betle* phytocompounds was predicted based on rule of five (Lipinsky *et al*., 2001). Solubility of drug was predicted by QP log S and Octanol/water partition coefficient (QP log Po/w) estimation. The brain / blood partition coefficient values - QP log BB (Luco and Kelder *et al*. 1999), Caco-2 cell permeability assay – QPP Caco (Yazdanian *et al*., 1998 and Irvine *et al*., 1999), MDCK (Madin-Darby Canine Kidney) cell permeability assay - QPP MDCK (Irvine *et al*., 1999), metabolic reactions prediction, IC 50 prediction for blockage of HERG K+channels (Cavalli *et al*. 2002 and De Ponti *et al*. (2002) and Skin permeability assay - QP logKp (Potts and Guy, 1992) were conducted.

## RESULTS and DISCUSSION

### Identification of bioactive compounds by Gas Chromatography – Mass Spectrometry (GC-MS) analysis

The phytocompounds detected in the GC-MS analysis of *P.betle* was shown in table 1 with their retention time (RT) and the area percentage. Also GC-MS chromatogram of alcoholic extract of *P. betle* extract was presented in figure 2. Totally 32 phytocompounds in *P.betle* was identified in the GC-MS analysis. The major phytocompounds were benzoic acid,3,5-dimethyl; 4-allyl-1,2-diacetoxy benzene (Acetoxy chavichol) and 4-carbamoyl-2,5-dimethyl pyridine (70.97 per cent). Benzene propanoic acid and 4-chromonol was the other major compounds (6.31 per cent) detected in analysis followed by gamma and beta sitosterol detected at 6.20 per cent. Also 2 methyl-Z,Z-3,13-octa decadienol and 1,2-15,16-diepoxy hexadecane were detected at 3.62 per cent and 3-Allyl-6-methoxyphenol (Chavibetol) and Eugenol were detected at 2.70 percent in the analysis.

**Figure 2.**
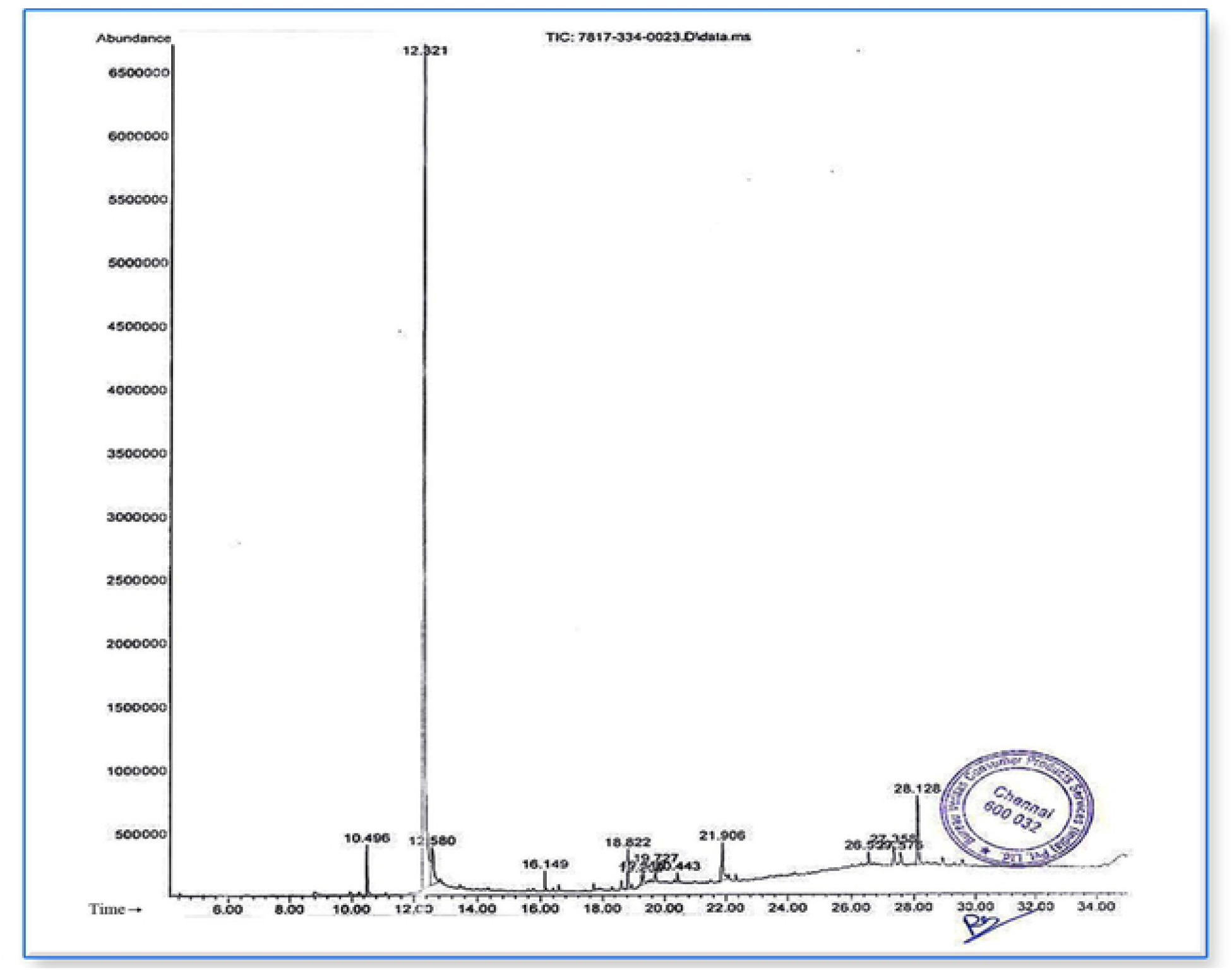
GC-MS chromatogram of Piper betle alcoholic extract.

**Table.1.** Phytocompounds detected in GC-MS analysis of *P.betle* alcoholic extract.

| **S. No.** | **Name of the Phytocompound** | **Retenti on time (minut es)** | **Area detected (percentage)** |
| --- | --- | --- | --- |
| 1 | a. 3-Allyl-6-methoxyphenol (Chavibetol)<br>b. Eugenol<br>c. Eugenol | 10.497 | 2.70 |
| 2 | a. Benzoic acid ,3,5-dimethyl<br>b. 4-Allyl-1, 2-diacetoxy benzene (Acetoxy chavicol)<br>c. 4 Carbamoyl – 2 - 5 dimethyl pyridine | 12.322 | 70.97 |
| 3 | a. Benzene propanoic acid , 2 - octyl ester<br>b. 4 Chromanol<br>c. Benzoic acid, 2,5-dimethyl | 12.578 | 6.31 |
| 4 | a. 9-Octadecyne<br>b. 6-Octen-1-ol, 3,7-dimethyl - propanaote<br>c. Bicyclo (3.1.1) heptane, 2,6,6-trimethyl | 16.151 | 0.93 |
| 5 | a. Phytol<br>b. Phytol<br>c. Phytol | 18.821 | 2.18 |
| 6 | a. Linoleic acid ethyl ester<br>b. (R)-(-)-14 Methyl- 8-hexadecyn1-ol<br>c. 9,12 Octadecadienoic acid (Z,Z) | 19.261 | 0.57 |
| 7 | a. 9,17-Octa decadienal, (Z)<br>b. 2-Methyl-Z,Z-3,13- octadecadienol<br>c. 9,12-Octadecadienoic acid (z,z) | 19.314 | 0.795 |
| 8 | a. Phytol, acetate<br>b. Cyclopropaneoctanal, 2- octyl<br>c. 11,13-Dimethyl -12-tetradecen-1-ol acetate | 19.725 | 1.18 |
| 9 | a. Cyclopropaneoctanal,<br>b. 2-Octyl-cis-13 –Octadecenoic acid<br>c. Z-B-Methyl-9-tetradecenoic acid | 20.444 | 0.60 |
| 10 | a. Cyclododecyne<br>b.2-Methyl-Z,Z-3,13- octadecadienol<br>c.1,2-15,16-Diepoxy hexadecane | 21.907 | 3.62 |
| 11 | a. Vitamin E,<br>b. Gamma tocopherol<br>c. Alpha tocopherol | 26.557 | 1.17 |
| 12 | a. Silicic acid, diethyl bis(trimethylsilyl) ester<br>b. Methyl-2-phenyllindolizine<br>c. Methyltris (trimethylsiloxy) silane | 27.359 | 1.71 |
| 13 | a. Methyltris (trimethylsiloxy) silane<br>b. Arsenous acid, tris (trimethylsilyl) ester<br>b. N-Methyl-1-adamantane acatamide | 27.573 | 1.08 |
| 14 | a. Gamma sitosterol<br>b. Beta sitosterol<br>c. Beta sitosterol | 28.126 | 6.20 |

### *In silico* analysis - Molecular docking study

The need for a rapid search for molecules that may bind to targets of biological interest is of crucial importance in the drug discovery process. One way of achieving this is through *in silico* analysis of molecules that contains relatively many hits against a particular target. Hence, in the present study the Schrodinger software was used to display computer generated interaction image of phytochemicals with target structure in a variety of orientations.

The Schrodinger module docked different conformational structures of *P.betle* phytocompounds and allopurinol into xanthine oxidase enzyme structure. The software was used to generate a pose after docking and the energetically most favourable pose was identified by GLIDE XP scoring. Scoring was done for all phytochemicals in the study, which were then ranked by their scores. Also the GLIDE energy, MM-GBSA energy, hydrogen bond and Pi-Pi interactions of phytochemicals with xanthine oxidase site were used to analyse the results in comparison with allopurinol. The results of docking studies of *P.betle* phytocompounds and allopurinol were presented in table 2 and 3. The parameters *viz*. GLIDE XP docking score (Kcal/mol), GLIDE energy (Kcal/mol), MM-GBSA energy (Kcal/mol) and the interaction with xanthine oxidase sites were computed and presented in the tables.

**Table 2.** Molecular docking of *P. betle* phytochemicals with xanthine oxidase.

| **S. No** | **Compound Name** | **GLIDE XP score (Kcal/mol)** | **GLIDE energy (Kcal/mol)** | **MM- GBSA (Kcal/mol)** | **Interaction sites** |
| --- | --- | --- | --- | --- | --- |
| 1. | 4-Chromanol | - 6.881 | - 25.427 | - 41.788 | GLH 802, PHE 1009, PHE 914 |
| 2. | 4-Allyl-1,2-diacetoxybenzene / Acetoxy chavichol | - 6.726 | - 42.822 | - 46.346 | SER 1080, SER1082, ALA 1079. |
| 3 | Eugenol | - 5.645 | - 24.813 | - 42.794 | GLH 802, PHE 1009, PHE 914 |
| 4 | Benzoic acid, 3, 5-dimethyl | - 5.367 | - 30.435 | - 36.978 | GLN 1040 |
| 5 | 3-Allyl-6-methoxy phenol / Chavibetol | - 5.342 | - 29.918 | - 35.529 | GLU 1261 |
| 6 | 2-Methyl-Z,Z-3,13-octadecadienol | - 5.377 | - 39.070 | - 75.694 | SER 1080 |
| 7 | 14-Methyl-8-hexadecen-1- ol.1 | - 5.330 | - 40.248 | - 54.697 | SER 1080, VAL 1259, GLY 1260. |
| 8 | Linolenic acid ethyl ester | - 5.312 | - 38.558 | - 40.389 | THR 1010, ARG 880. |
| 9 | Gamma tocopherol | - 5.261 | - 36.988 | - 37.540 | GLN 1194 |
| 10 | 1,2-15,16 -Diepoxy hexa decane | - 5.216 | - 39.239 | - 64.231 | VAL 1011 |
| 11 | Bicyclo heptane, 2,6,6 - trimethyl | - 4.831 | - 24.891 | - 29.326 | No interaction |
| 12 | 6-Octen-1-ol,3,7-dimethyl propanaote | - 4.766 | - 36.970 | - 42.321 | GLN 1040 |
| 13 | Phytol | - 3.895 | - 39.594 | - 73.553 | GLN 1194 |
| 14 | 9,12-Octadecadienoic acid | - 3.244 | - 40.826 | - 39.862 | GLY 1260, VAL 1259, THR 1083. |
| 15 | Methyl - 2 Phenyl lindolizine | - 3.190 | - 30.824 | - 40.025 | No interaction |
| 16 | Silicic acid,diethylbis(trimethylsilyl) ester | - 2.613 | - 27.606 | -19.943 | No interaction |
| 17 | 9,17 - Octa decadienal | - 1.773 | - 37.019 | - 35.943 | No interaction |
| 18 | Cyclopropane octanal | - 1.417 | - 36.706 | - 35.408 | No interaction |

**Table 3.** Molecular docking of Allopurinol with xanthine oxidase.

| **S.**<br>**No** | **Compound Name** | **GLIDE XP score**<br>**(Kcal/mol)** | **GLIDE energy**<br>**(Kcal/mol)** | **MM- GBSA**<br>**(Kcal/mol)** | **Interaction sites** |
| --- | --- | --- | --- | --- | --- |
| 1 | Allopurinol | - 4.535 | - 32.676 | - 30.816 | THR 1010, ARG 880,<br>GLH 802, PHE 1009, PHE 914. |

The standard drug allopurinol showed GLIDE XP docking score of - 4.535 Kcal/mol and the GLIDE energy of - 32.676 Kcal/mol and the MM - GBSA bind energy of - 30.816 Kcal/mol. The allopurinol had three hydrogen bond interactions (THR 1010, ARG 880, GLH 802 sites of xanthine oxidase) and two Pi - Pi interactions (PHE 1009, PHE 914 sites of xanthine oxidase) as shown in Figure 3. The standard drug allopurinol docking results are in agreement with Umamaheswari *et al*. (2011).

**Figure 3.**
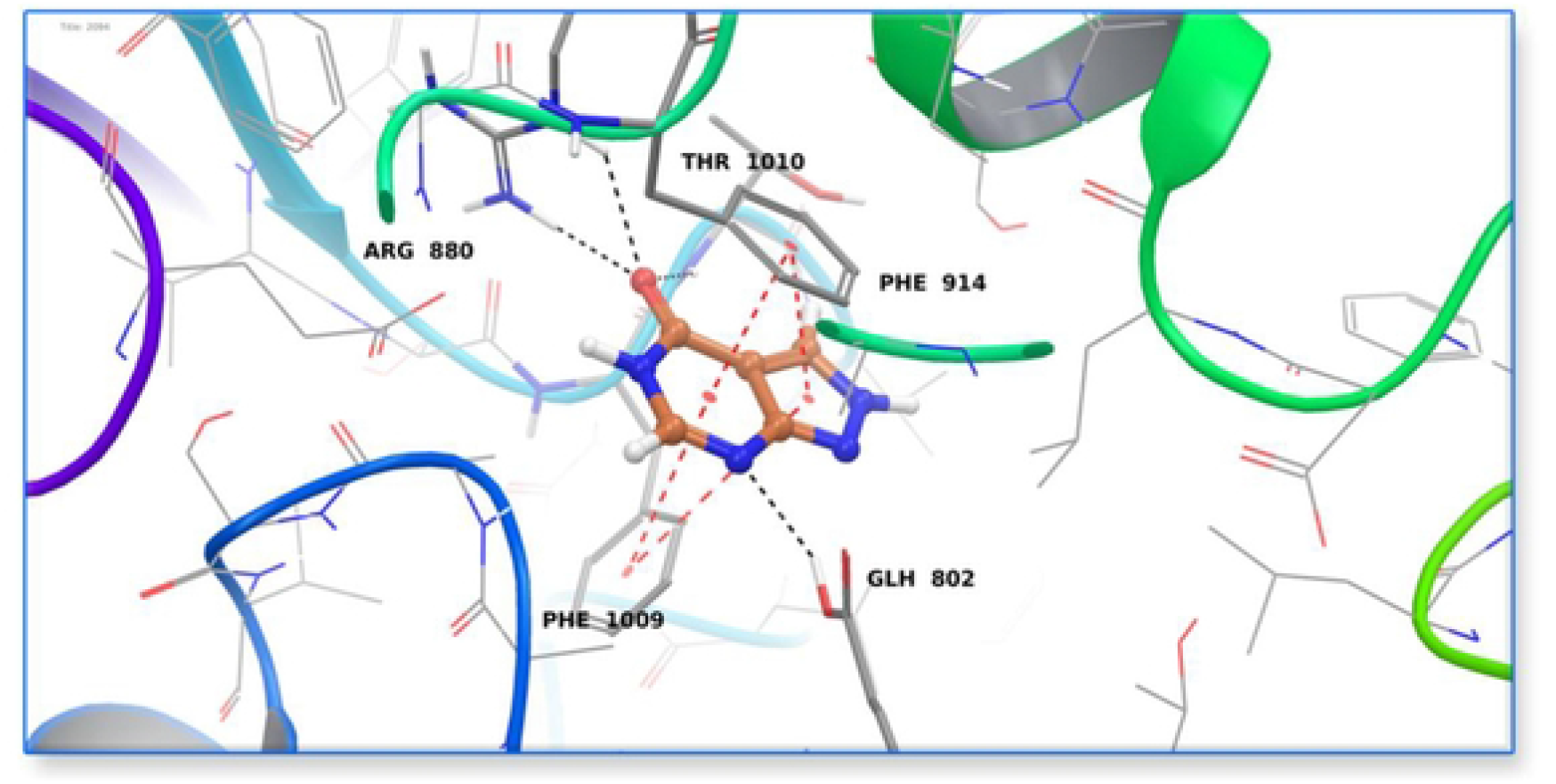
In silico Study – Allopurinol docking with Xanthine oxidase - 3D image.

Among the 32 phytocompounds of *P.betle* eighteen had favourable binding affinity with xanthine oxidase target and they were docked. The *P.betle* phytochemicals GLIDE XP docking score were found to be ranging between - 1.417 Kcal/mol (cyclopropane octanal) and - 6.881 Kcal/mol (4 chromonal). The 12 phyto compounds of *P. betle* viz. 4-chromanol (−6.881 Kcal/mol), 4-allyl-1,2-diacetoxybenzene (−6.726 Kcal/mol), eugenol (−5.645 Kcal/mol), benzoic acid,3,5-dimethyl (−5.367 Kcal/mol), 3-allyl-6-methoxy phenol (−5.342 Kcal/mol), 2-methyl-Z,Z-3,13-octadecadienol (−5.377 Kcal/mol), 14-methyl-8-hexadecen-1- ol.1 (−5.330 Kcal/mol), Linolenic acid ethyl ester (−5.312 Kcal/mol), gamma tocopherol (−5.261 Kcal/mol), 1,2-15,16-diepoxy hexa decane (−5.216 Kcal/mol), bicyclo heptane,2,6,6-trimethyl (−4.831 Kcal/mol) and 6-octen-1-ol,3, 7-dimethyl-propanaote (−4.766 Kcal/mol) had higher docking score when compared to the standard drug allopurinol (−4.535 Kcal/mol) and the remaining 6 compounds of *P.betle* showed lesser docking score than allopurinol, since they did not have hydrogen bond and Pi-Pi interactions.

The *P.betle* phytocompounds showed GLIDE energy ranging between - 24.813 Kcal/mol (eugenol) and - 42.822 Kcal/mol (4-allyl-1, 2-diacetoxy benzene). Totally 11 phytocompounds of *P.betle* viz. 4-allyl −1 −2 - diacetoxy benzene (−42.822 Kcal/mol), 9,12 octadecadienoic acid (−40.826 Kcal/mol), 14-methyl-8-hexadecen-1-ol.1 (−40.248 Kcal/mol), phytol (−39.594 Kcal/mol), 1,2-15,16-diepoxyhexadecane (−39.239 Kcal/mol), 2-methyl- Z,Z-3,13-octadeca dienol (−39.07 Kcal/mol), linolenic acid ethyl ester (−38.558 Kcal/mol), 9,17-octadecadienal (−37.019 Kcal/mol), gamma tocopherol (−36.988 Kcal/mol), 6-octen-1-ol,3,7-dimethyl propanaote (−36.97 Kcal/mol) and cyclopropane octanal (−36.706 Kcal/mol) showed higher GLIDE energy than standard allopurinol (−32.676 Kcal/mol) and the remaining seven compounds showed lesser GLIDE energy than allopurinol.

The GLIDE score and GLIDE energy values were used to identify the prime phytocompounds having xanthine oxidase inhibitory activity. The higher negative energy values represented high binding affinity between the receptor and ligand molecules and indicated higher efficiency of the bioactive compounds (Mani *et al*., 2016). The results of the present study revealed that the acetoxy chavichol compound of *P.betle* had high binding affinity with xanthine oxidase enzyme than the standard drug allopurinol.

The MM-GBSA energy value of *P.betle* phytochemicals ranged between - 19.943 Kcal/mol (Silicic acid, diethyl bis (trimethylsilyl) ester) and - 75.694 Kcal/mol (2-methyl-Z,Z-3,13-octadecadienol). Except silicic acid, diethyl bis (trimethyl silyl) ester and bicyclo heptane, 2,6,6-trimethyl (−29.326 Kcal/mol), all the *P.betle* phytocompounds showed higher MM - GBSA energy value than standard drug allopurinol (−30.816 Kcal/mol). The MM-GBSA value generally showed free energy of binding which is an effective way to measure binding strength (Genheden and Ryde, 2015). As per the results of present study, the acetoxy chavichol compound in *P.betle* showed higher MM-GBSA value than the standard drug allopurinol and showed the better binding strength of docked complex.

Chromonal, the topmost interacting compound of *P. betle* had three interactions with xanthine oxidase sites of which one hydrogen bond interaction with the amino acid GLH 802 and two Pi - Pi interactions with PHE 1009 and PHE 914 aminoacids as shown in figure 4. The other *P.betle* phytochemicals *viz*., Acetoxy chavichol, Eugenol and Chavibetol interactions with xanthine oxidase sites were shown in figure 5-7.

**Figure 4.**
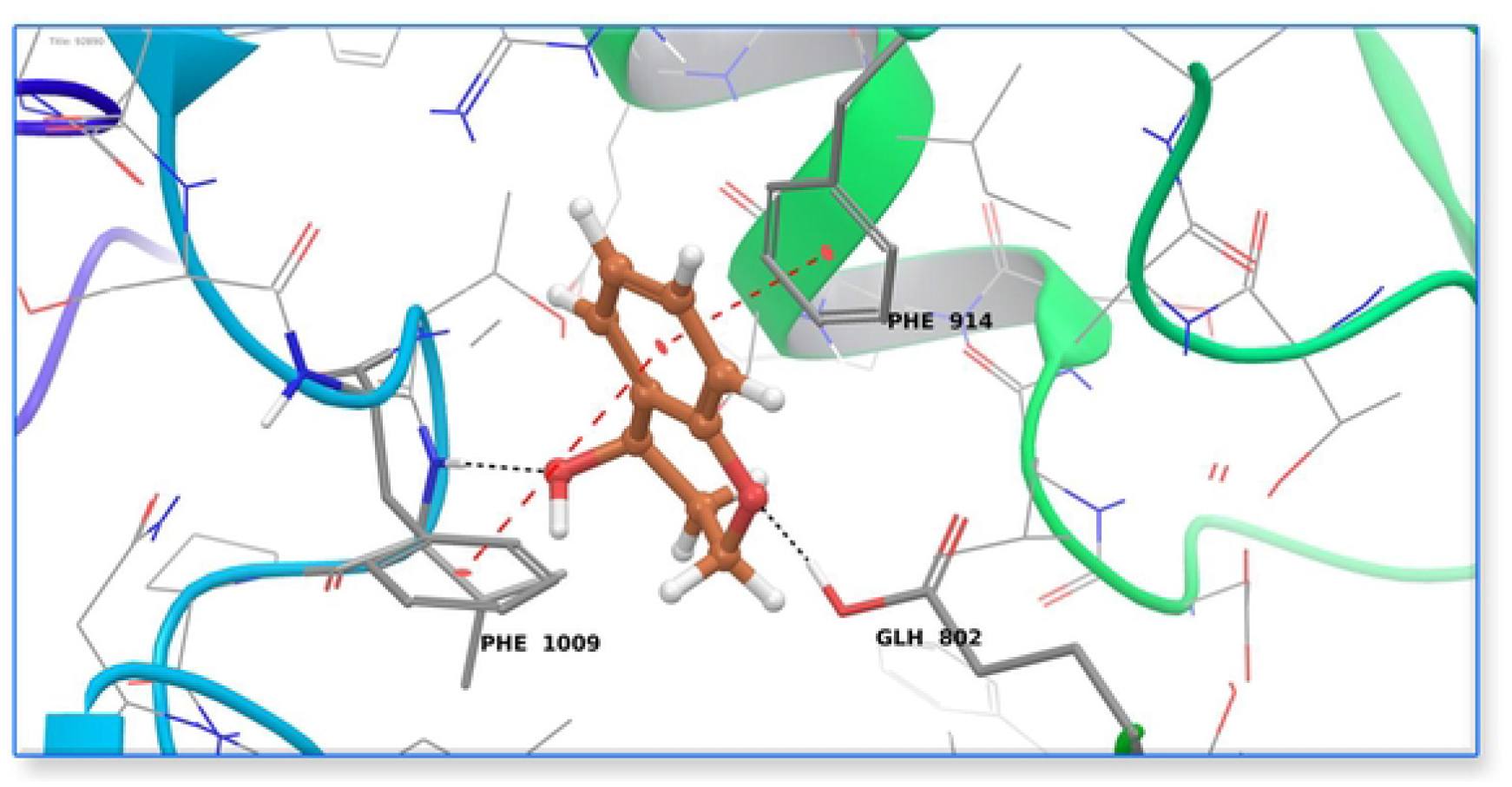
In silico Study – Chromonol (Phytocompound of P.betle) docking with Xanthine oxidase - 3D image.

**Figure 5.**
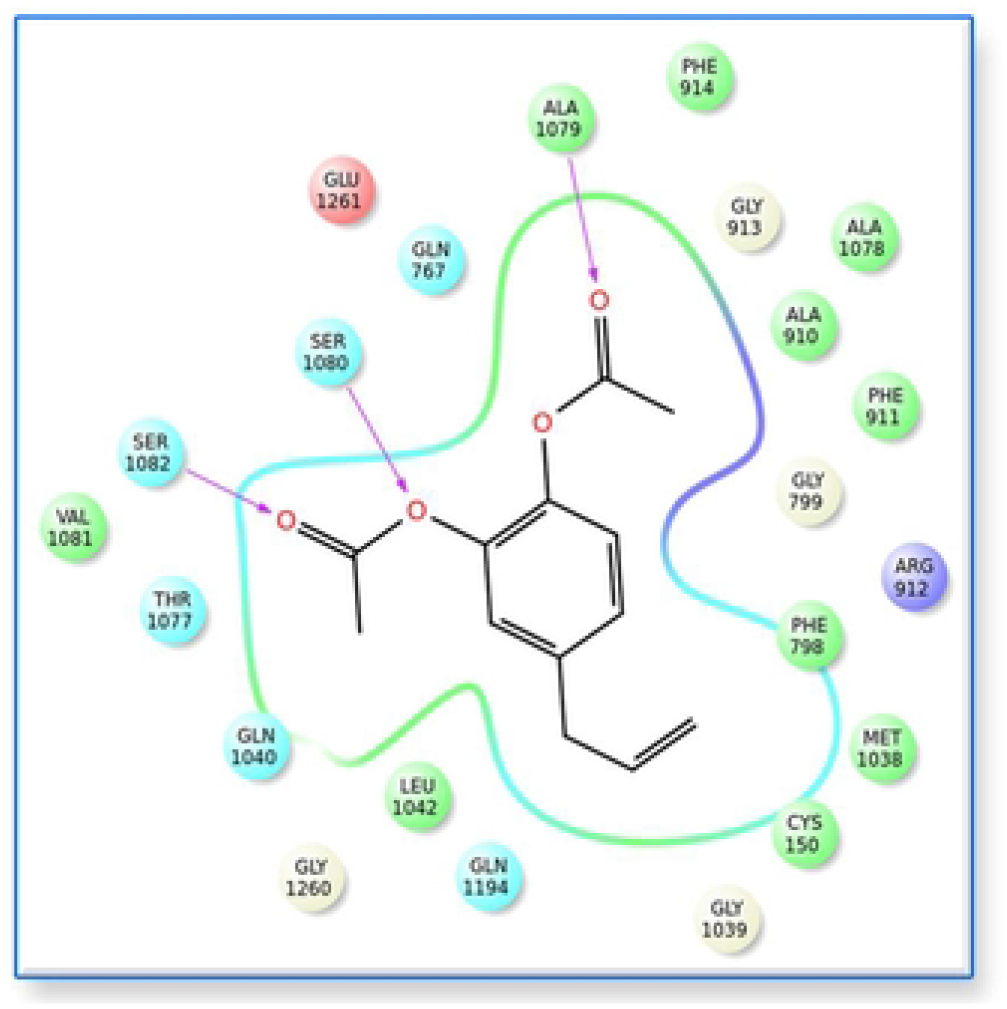
In silico Study – Acetoxy Chavichol (Major phytocompound of P.betle) docking with Xanthine oxidase - 2D image.

**Figure 6.**
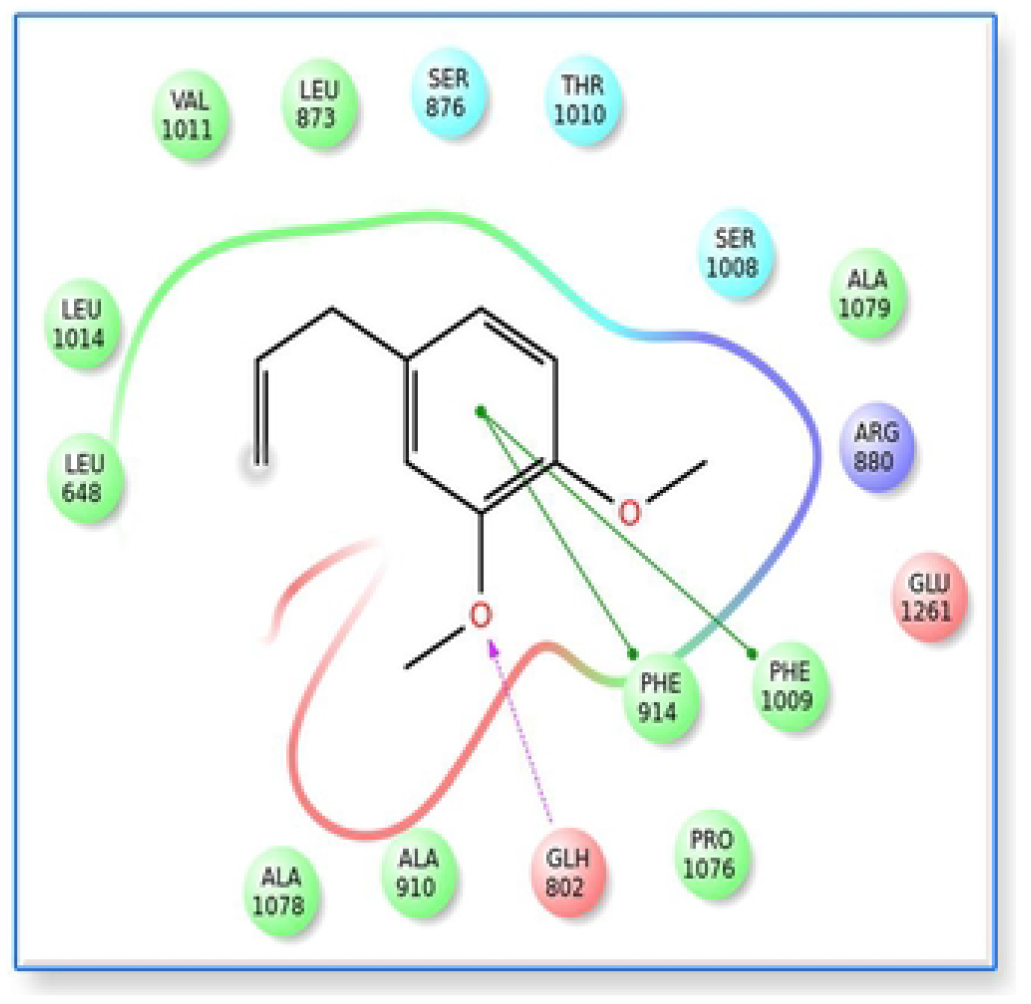
In silico Study – Eugenol (Phytocompound of P.betle) docking with Xanthine oxidase - 2D image.

**Figure 7.**
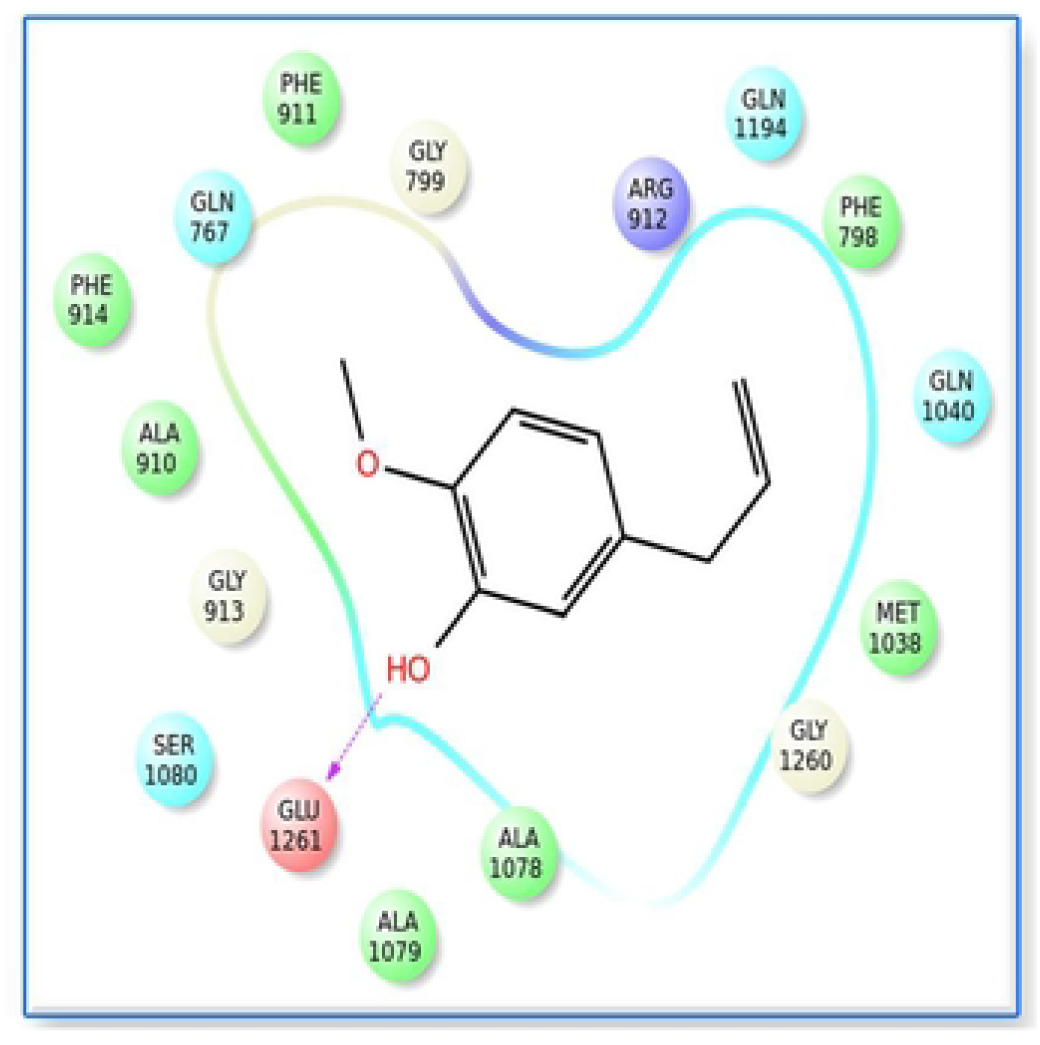
In silico Study – Chavibetol (Phytocompound of P.betle) docking with Xanthine oxidase - 2D image.

Chromonol and Eugenol were the most potential compounds of *P.betle* and they showed both hydrogen bond and Pi-Pi interactions with the target xanthine oxidase as that of standard drug allopurinol. In general, the presence of both hydrogen bond and Pi-Pi hydrophobic interactions between the phytocompound and the target were found to be responsible for the antigout / antihyperuricimic activity (Umamaheswari *et al*., 2011). Hence presence of both hydrogen bond and Pi-Pi interactions in the *P.betle* phytocompounds might be expected to produce antihyperuricimic activity in *invivo* study also.

### ADMET analysis

In this study, the ADMET properties of six phytocompounds of *P.betle* were analysed using Qik Prop tool. The recommended values for all parameters and the molecular weight, no.of violations in rule of five, percentage of oral absorption and other kinetic parameter and toxicity profile results were shown in the table 4.

**Table. 4.** Prediction of Absorption, distribution, metabolism, excretion and toxicity profile of *P.betle* phytocompounds using QikProp module of Schrödinger.

| Compound Name | Molecular Weight | Rule of five | Percentage of oral absorption | QP logS | Qp log po/w | Qp log kp | Qp log BB | Qpp Caco | QPP MDCK | No.of metabolic reactions | QPlog HERG |
| --- | --- | --- | --- | --- | --- | --- | --- | --- | --- | --- | --- |
| Recommended Values | 130.0 – 725.0 | Violations is maximum is 4 | > 80% is high<br>< 25% is poor | – 6.5 to 0.5 | –2.0 to 6.5 | – 8.0 to –1.0 | –3.0 to 1.2 | < 25 – poor,<br>> 500 - great | < 25 – poor,<br>> 500 - great | 1-8 | below –5 |
| 4 - Chromanal | 150.1 | 0 | 100.00 | -1.549 | 1.558 | -1.593 | 0.184 | 3918.257 | 2164.7 | 5.53 | -3.310 |
| 4-Allyl-1,2-diacetoxybenzene / Acetoxy chavichol | 178.2 | 0 | 100.00 | -3.657 | 2.816 | -0.695 | 0.090 | 9906.0 | 5899.2 | 4.55 | -3.713 |
| 3-Allyl-6-methoxy phenol / Chavibetol | 164.2 | 0 | 100.00 | -2.432 | 2.678 | -1.603 | -0.133 | 3047.2 | 1649.6 | 5.00 | -4.036 |
| Methyl octadecadienol | 280.493 | 1 | 100.00 | -6.067 | 6.098 | -0.475 | -0.647 | 5328.2 | 3017.9 | 4.40 | -5.177 |
| Eugenol | 164.2 | 0 | 100.00 | -2.250 | 2.683 | -1.381 | -0.020 | 3976.3 | 2199.4 | 4.10 | -3.833 |
| Benzoic acid 3,3 dimethyl | 150.1 | 0 | 81.015 | -1.750 | 1.887 | -3.038 | -0.352 | 253.399 | 142.683 | 3.20 | -1.703 |

**Qp log S-** Predicted aqueous solubility, **Qp logP o/w-** Predicted octanol/water partition coefficient, **Qp log kp -** Predicted skin permeability , **Qp log BB -** Predicted brain/blood partition coefficient, **Qpp caco -** Predicted apparent Caco-2 cell permeability in nm/sec, **QPP MDCK-** Predicted apparent MDCK cell permeability in nm/sec, **Qplog HERG-** Predicted IC50 value for blockage of HERG K<sup>+</sup> channels.

Based on the induced fit docking analysis, six molecules which showed better GLIDE score and better energy values than allopurinol were selected and used for ADMET analysis. Since 60% of drugs fail to attain pharmacokinetic properties during drug development (Nisha et al., 2016), the ADMET analysis could help in the earlier prediction of pharmacokinetic properties and it reduces the late stage compound attrition during drug discovery process thereby reduces the economic burden in drug development research.

QikProp tool was used in the present study and it generates set of physicochemically significant descriptors which further evaluates ADMET properties. Lipinski’s rule of 5 is a rule of thumb to evaluate drug-likeness, or determine if a chemical compound with a certain pharmacological or biological activity has properties that would make it a likely orally active drug in humans. The rule describes molecular properties important for a drug’s pharmacokinetics in the human body, including its ADME. This parameter determines the number of property descriptors calculated via QikProp which fall outside from the optimum range of values for 95% of noted drugs. In the present study, the rules like, molecular weight < 500, QP log P o/w < 5, donor HB ≤ 5, acceptor HB ≤ 10 were calculated for each phytocompound. Except Methyl octadecadienol, none of selected phytocompounds showed violations of rules which reflected the drug likeness of the compounds of *P.betle*. The molecular weight of the phytocompounds was within the prescribed limit. Hence all of these compounds could expect to have good absorption and excretion in the system. Also, high percentage of oral absorption rate was predicted for all compounds.

The solubility in the intestinal fluid is an important property of oral drugs since insufficient solubility may limit the intestinal absorption through the portal vein system to obtain a therapeutic effect when systemic effects are warranted. Water-soluble compounds greatly facilitate many drug development activities, primarily because of the ease of handling and formulation. For oral administration, solubility is a major property influencing absorption. Similarly, a drug meant for parenteral usage has to be highly soluble in water to deliver a sufficient quantity of the active ingredient (Daina *et al*., 2017). QP log S value showed the solubility of compounds in water (log S). The water solubility (QP log S) is critical for absorption and distribution of drugs within the body and it ranged between - 6.067 to - 1.549 and all the readings were within the recommended range which showed the possible good absorption and distribution of *P.betle* phytocompounds.

Likewise, the partition coefficient between octanol and water (log P o/w) is a common descriptor to measure lipophilicity. Lipophilicity of the compounds is related to the permeability through biological membranes. It could be decreased when lipophilicity is too low, whereas very hydrophilic compounds are usually not able to diffuse passively through them (Lagorce, *et al*., 2017). The partition coefficient (QP log P o/w) is also critical for absorption and distribution of drugs within the body and it ranged between 1.558 to 6.098 and all the readings were within the recommended range which again confirms the good absorption and distribution of *P.betle* phytocompounds.

The permeability across the skins and blood-brain barrier is a crucial factors determining effectiveness of drugs in the dermal and CNS systems. The blood/brain partition coefficients (log B/B) were computed in the present study and used as a predictor for access to the central nervous system (CNS). The skin permeability factor (log Kp) and brain/blood partition coefficient values (log BB) values were ranged in between – 3.038 to - 0.475 and – 0.647 to 0.184, respectively and they were within the recommended values which predicted the excellent permeability of *P.betle* phytocompounds across skin and central nervous system.

Caco2 cell permeability assay was performed to know about the phytocompound permeability in gut-blood barrier. This parameter is a key factor governing drug metabolism and its access to biological membranes, ranged from 253.399 to 9906 in the present study which validates the greater permeability of all the compounds in gut-blood barrier except Benzoic acid 3,3 dimethyl molecule.

The blood-brain barrier permeability prediction study was conducted using MDCK (Madin-Darby Canine Kidney) cell permeability assay. MDCK assay are widely used to make oral absorption estimates, since the cells also express transporter proteins, but only express very low levels of metabolizing enzymes (Veber et al. 2002). The predicted values were ranges from 142.683 to 5899.20 which validated the greater blood-brain barrier permeability of all the compounds except Benzoic acid 3,3 dimethyl molecule.

An estimated number of possible metabolic reactions has also been predicted by QikProp and used to determine whether the molecules can easily gain access to the target site after entering the blood stream. In the present study the metabolic reactions of all the compounds are within the prescribed limit.

Human ether-a-go-go related gene (HERG) encodes a potassium ion (K+) channel that is implicated in the fatal arrhythmia known as torsade de pointes or the long QT syndrome (Hedley *et al*. 2009). The HERG K+ channel, which is best known for its contribution to the electrical activity of the heart that coordinates the heart’s beating, appears to be the molecular target responsible for the cardiac toxicity of a wide range of therapeutic drugs (Vandenberg 2001). Many drugs have been withdrawn from the market because of the serious hERG-related cardiotoxicity. HERG has also been associated with modulating the functions of some cells of the nervous system and with establishing and maintaining cancer like features in leukemic cells (Chiesa *et al*. 1997). Thus, HERG K+ channel blockers are potentially toxic and the predicted IC50 values often provide reasonable predictions for cardiac toxicity of drugs in the early stages of drug discovery (Aronov, 2005). The recommended range for predicted log IC50 values for blockage of HERG K+ channels (logHERG) is > - 5. The range of values predicted in the analysis was in-between - 1.703 to to - 5.177 which revealed that all the compounds are safer to use, except Methyl octadecadienol.

## Conclusion

*Insilico* study revealed that the GLIDE XP score, GLIDE energy and MM-GBSA energy values revealed that 12 phytocompounds of *P.betle* had superior scoring than allopurinol and 11 phytocompounds of *P.betle* had high negative GLIDE energy value than allopurinol. Also, 16 phytocompounds of *P.betle* showed high MM-GBSA energy value than allopurinol. Among the different phytocompounds, the acetoxychavichol, chromonol and eugenol are the most promosing phytocompounds present in the *P.betle* herb and they showed higher GLIDE XP score, GLIDE energy and MM – GBSA energy and had favourable interactions with xanthine oxidase enzyme.

Further, the drug likeness revealed by the all six tested *P.betle* phytocompounds and many of the pharmacokinetic parameter values of *P.betle* compounds were within the acceptable range defined for human use, during ADMET analysis which indicates their potential as drug-like molecules. Hence it was concluded that the *insilico* and ADMET profile study on the phytocompounds of *P.betle* predicted their promising potential xanthine oxidase inhibition activity and they could be developed as alternate molecules against synthetic agents by *invivo* experiments for the treatment of xanthine oxidase associated diseases like hyperuricimia and cardiovascular disorders.

## Acknowledgment

The authors wish to thank the authorities of Tamil Nadu Veterinary and Animal Sciences University, Chennai, Tamil Nadu, India and Alagappa university, Karaikudi, Tamil Nadu India for the grant of permission to conduct the study.

